# A coupled fluorescence assay for high-throughput screening of polyurethane-degrading enzymes

**DOI:** 10.1101/2025.07.09.663332

**Authors:** Liujun Yang, Xianjun Deng, Renju Liu, Jianyang Li, Junyao Zhai, Xiaoxiao Ma, Arthur K.L. Lin, Jiguang Wang, Zongze Shao, Hongyuan Lu

**Affiliations:** Thrust of Bioscience and Biomedical Engineering, The Hong Kong University of Science and Technology (Guangzhou), Guangzhou 511400, China; Third Institute of Oceanography, Ministry of Natural Resources, Xiamen 361005, China; Practice in Engineering Education, The Hong Kong University of Science and Technology (Guangzhou), Guangzhou 511400, China; Division of Life Science, Department of Chemical and Biological Engineering, The Hong Kong University of Science and Technology, Clear Water Bay, Kowloon, Hong Kong SAR, China

**Keywords:** fluorescence, high-throughput screening, polyurethane, enzyme engineering, biodegradation, ethylene glycol

## Abstract

The global accumulation of plastic waste, particularly persistent polymers like polyurethane (PU), demands urgent solutions. Enzymatic depolymerization offers a viable strategy for PU waste valorization. However, progress has been hindered by the lack of reliable high-throughput screening (HTS) assays capable of precise and quantitative evaluation of enzymatic activity. Current enzyme screening assays face significant limitations due to poor substrate relevance and inadequate quantification methods. Here, we present a novel HTS assay featuring a chemically defined, synthetic poly (ethylene adipate)-based PU substrate that enables unambiguous structure-activity analysis and direct quantification of degradation products (adipic acid and ethylene glycol). Coupled with this substrate is a highly sensitive fluorescence-based detection cascade, in which released ethylene glycol is stoichiometrically converted to resorufin via a two-enzyme system (glycerol dehydrogenase and diaphorase). This assay overcomes key limitations of commercial substrates (e.g., Impranil DLN) by providing quantitative, real-time monitoring of PU hydrolysis with molecular precision. Validation with known PU-degrading enzymes demonstrated that the assay can sensitively distinguish differences in enzymatic activity with high reproducibility and quantitative accuracy. Our platform enables rapid, cost-effective screening of enzyme libraries and engineered variants, significantly advancing enzyme discovery and optimization efforts. By bridging the gap between laboratory research and industrial application, this HTS assay accelerates the development of sustainable PU recycling solutions, offering a critical tool against plastic pollution.

**Graphical Abstract Legend:** **(A)** Workflow of the enzymatic degradation assay. Enzymes are incubated with chemically defined PEA-PU films. The resulting hydrolysate is collected and analysed using a fluorescence-based assay in microplate format (Ex/Em = 535/588 nm). **(B)** Fluorescence-based detection mechanism. Assay principle based on ethylene glycol (EG), one of the major degradation products. EG is converted through a two-step enzymatic cascade involving glycerol dehydrogenase (GldA) and diaphorase, leading to the generation of fluorescent resorufin from resazurin.

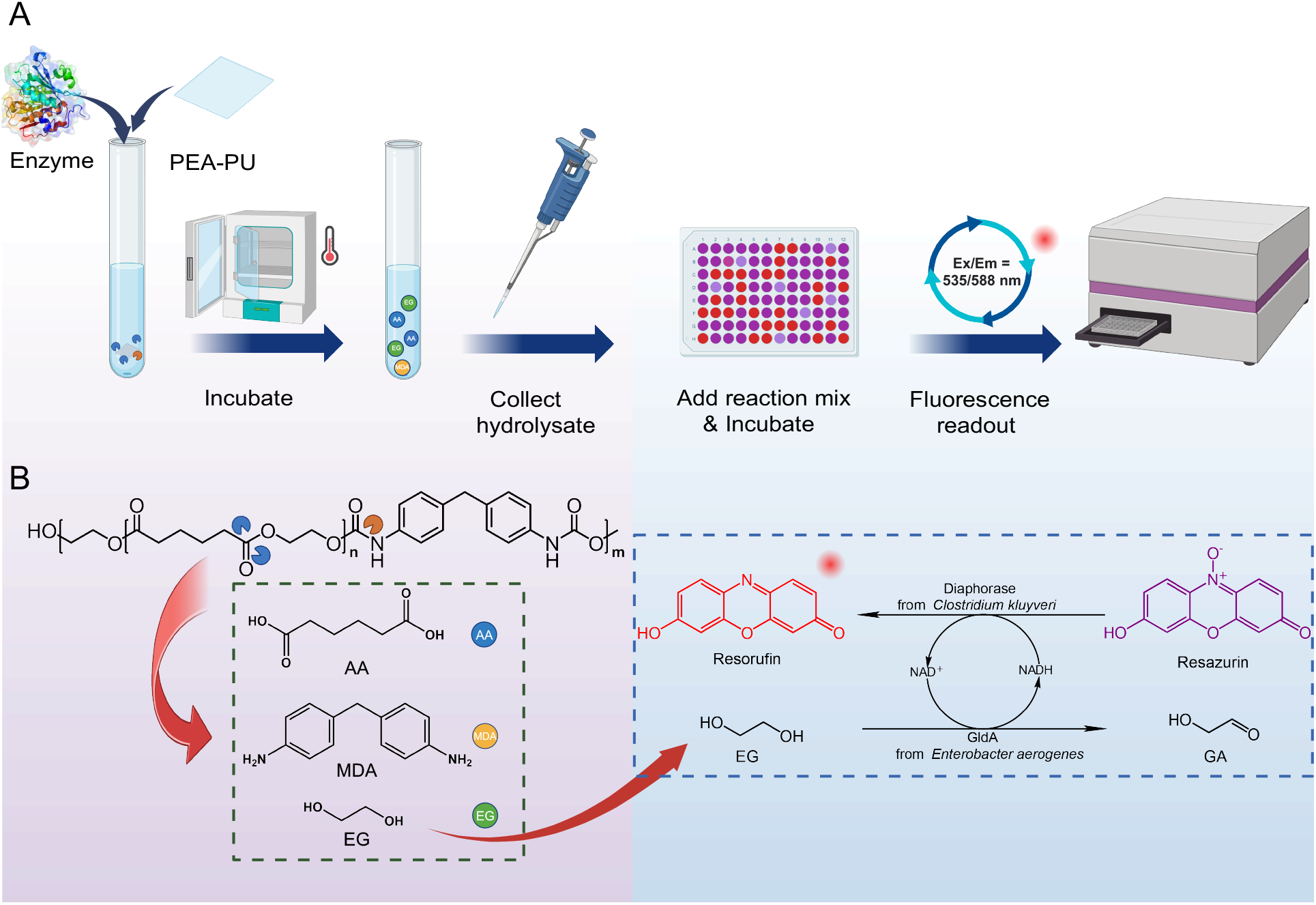

## 1. Introduction

Plastic pollution poses a pressing global challenge, threatening ecosystems ^1-3^ and human health ^4–9^. A major contributor to this crisis is polyurethane (PU), a synthetic polymer extensively used in foams (e.g., mattresses, insulation), elastomers (e.g., shoe soles, medical devices), and durable goods (e.g., automotive parts, synthetic leather) ^10–12^. Given the low recycling rate of PU waste ^12,13^, its environmental persistence highlights the urgent need for sustainable solutions ^13–15^.

Enzymatic depolymerization has emerged as a promising strategy for PU waste valorization, attracting growing research interest ^10,13–16^. However, progress in this field remains limited. This is largely due to persistent challenges in identifying, characterizing, and engineering enzymes that can efficiently degrade PU under industrial conditions ^14–18^.

A major bottleneck lies in current enzyme screening assays, which suffer from limitations in predictive accuracy and practical utility (Table 1). Conventional assays using low-molecular-weight model substrates (e.g., p-nitrophenyl esters or carbamates) often fail to predict enzyme activity against authentic polymeric PU, leading to false positives that misdirect enzyme discovery efforts ^14,15,16,19^. In contrast, while industrial PU materials (e.g., foams, films, elastomers) offer greater physiological relevance, their heterogeneity and slow degradation kinetics render them unsuitable for high-throughput screening (HTS) applications ^15,18,20,21^.

**Table 1:** Critical limitations of current assay formats for polyurethane (PU) biodegradation. A Comparative overview of model substrates, colloidal PU dispersions, and commercial PU materials used in PU degradation assays.

| Feature | LMW Model Compounds | PU Dispersions | Commercial PU Materials |
| --- | --- | --- | --- |
| Representative Examples | *p*-Nitrophenyl esters, carbamates | Impranil® DLN, Bayhydrol® 110 | PU foam, film, elastomer |
| Chemical Definition | Defined, synthetic molecules | Proprietary formulations | undefined mixtures or composites |
| Physiological Relevance | Low | Moderate | High |
| Readout Modality | Colorimetric absorbance | Turbidity clearing, halo formation | Weight loss, SEM, product analysis (e.g., HPLC) |
| Data Quality | Quantitative (model-limited) | Non-quantitative | Quantitative (method-dependent) |
| Screening Throughput | High | Moderate | Low |
| Bond-Type Specificity | Restricted to designed motifs | Ester-biased; limited urethane sensitivity | Broad; structurally diffuse |
| Mechanistic Insight | Indirect; lacks polymer context | Partial; limited structural resolution | Direct, but labor-intensive |

Consequently, many studies have turned to colloidal PU dispersions (e.g., Impranil DLN or Bayhydrol), accessing enzyme activity via clearing zone formation ^22^ or by monitoring OD reduction in PU emulsions upon enzyme or microbial addition ^23^. However, these proprietary mixtures are chemically undefined and only detect bulk esterase activity. These limitations hinder rigorous quantitative analysis, which is crucial for enzyme discovery and mechanistic studies ^14,15,20,24^.

To overcome these limitations, we developed an HTS assay using a synthetic PU substrate designed to release detectable degradation products upon enzymatic hydrolysis. By coupling this reaction to a detection cascade— where degradation products are enzymatically converted into an optical signal—we enable real-time kinetic measurements and direct activity comparisons.

This assay resolves key ambiguities in existing methods by delivering a quantitative, high-throughput measurement of PU hydrolysis kinetics, enabling rapid and precise enzyme variants screening. Rigorous validation with established PU-degrading enzymes confirms its high sensitivity, reproducibility, and robustness. Beyond serving as a powerful tool for enzyme discovery and engineering, this assay also establishes a foundation for mechanistic studies of PU biodegradation.

## 2. Results and Discussion

### 2.1. Synthesis and characterization of PEA-PU

To develop a quantitative assay for PU-degrading enzymes, we designed a novel polymeric substrate that is chemically well-defined and readily synthesizable. The polymer features a soft segment composed of poly (ethylene adipate) (PEA). Hydrolysis of its ester bonds produces two well-defined products: ethylene glycol (EG) and adipic acid (AA). Both can be easily quantified with high reliability. Specifically, the PEA-PU polymer was synthesized via a one-step, solvent-based step-growth polymerization ^23,25^ (Fig. 1A). The resulting polymer solution was then cast into films exhibiting high optical transparency and uniformity (Fig. 1B). This optical quality reflects excellent macroscopic homogeneity, which is desirable for consistent enzyme-substrate interactions in replicate assays.

**Figure 1:**
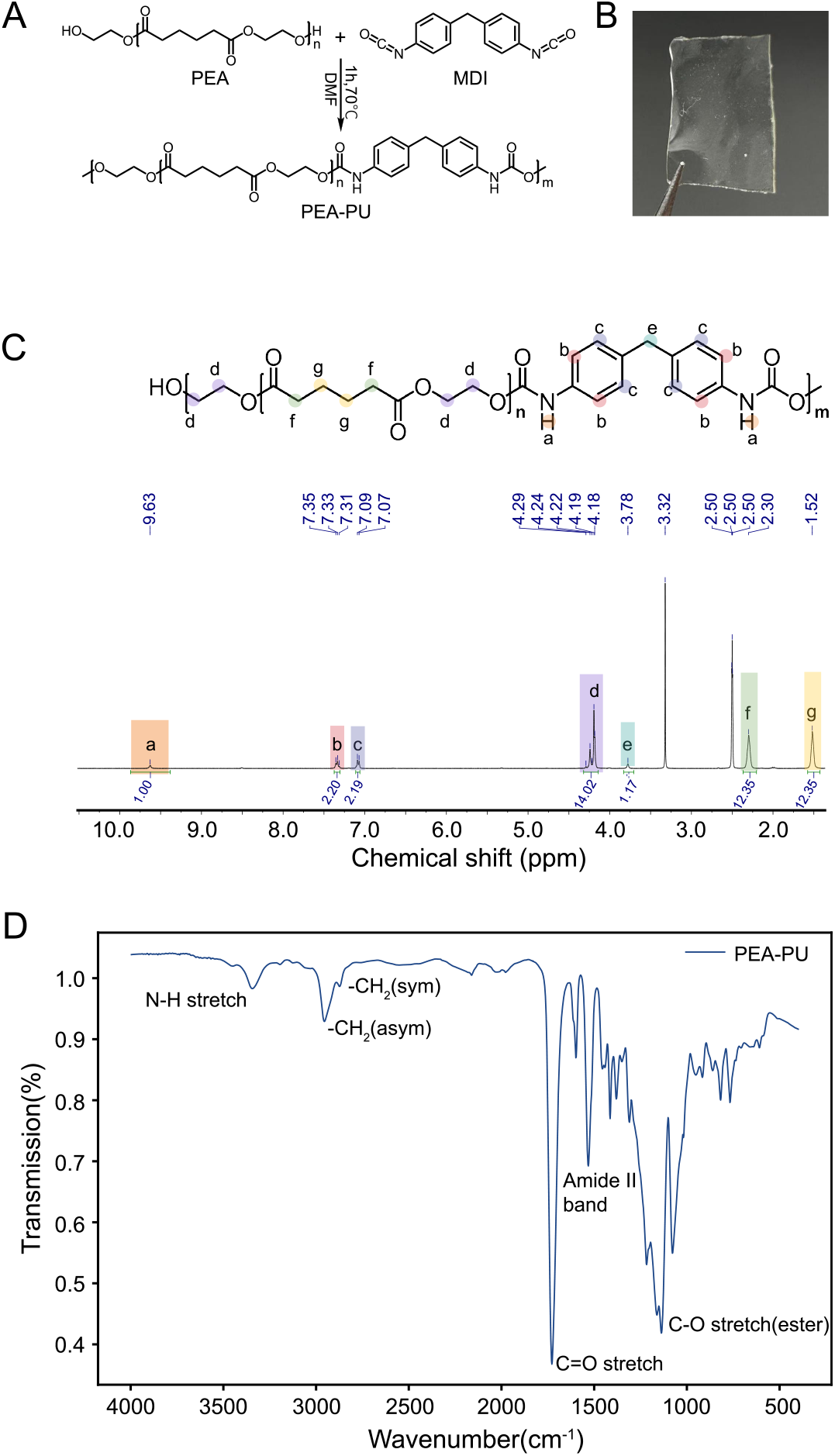
Synthesis and characterization of PEA-PU. **(A) Synthesis of PEA-PU via step-growth polymerization**. PEA-PU was synthesized by reacting pre-dried polyethylene adipate (PEA, Mn ≈ 1000 g/mol) with 4,4′-methylenebis (phenyl isocyanate) (MDI) at an NCO: OH molar ratio of 1:1 in anhydrous DMF at 70 °C under a nitrogen atmosphere for 1 hour. **(B) PEA-PU film**. The polymer solution was cast into a PTFE mold and dried under vacuum at 60 °C for 24 hours, yielding a film approximately 0.2 mm thick. **(C)** ^**1**^**H-NMR spectrum of PEA-PU**. δ 9.63 (br s, NH, urethane); 7.33 (t, J ≈ 8.6 Hz, 2 H) & 7.08 (d, J ≈ 8.2 Hz, 3 H) = aromatic H of MDI; 4.40–4.15 (m, ≈14 H) = overlapped O-CH_2_-CH_2_-O and α-CH_2_ next to carbonyl/urethane O; 3.78 (br s, Ar-CH_2_-Ar bridge, down-field half; up-field partner at ∼3.3 ppm is buried in the 2.6–2.3 ppm envelope); 2.60–2.35 (br m) = overlapped Ar-CH_2_-Ar partner + urethane α-CH_2_ ± minor end groups; 2.30 (br s) = adipate α-CH_2_; 1.52 (br s) = internal aliphatic CH_2_. **(D) FTIR spectrum of PEA-PU**.The spectrum exhibits characteristic absorption bands, including a composite carbonyl band at 1725 cm^−1^ (overlapping ester and urethane C=O stretches), a urethane N–H stretch at 3320 cm^−1^, an amide II band at 1530 cm^−1^ (N–H bending coupled with C–N stretching), and an amide III/ester C–O–C band at 1240 cm^−1^, collectively validating the expected urethane and ester functionalities in the polymer backbone.

To confirm the polymer’s chemical composition, we performed ^1^H NMR analysis. The ^1^H NMR spectrum (Fig. 1B) exhibited three diagnostic regions that unambiguously confirmed the polyester–urethane backbone architecture: (i) the urethane N–H proton at δ 9.63 ppm; (ii) aromatic and bridging methylene protons from the MDI-based hard segment at δ 7.07–7.35 and 3.78 ppm, respectively; (iii) O-linked methylene protons from the ethylene glycol units at δ 4.18–4.29 ppm, and aliphatic methylene protons from the adipate segment at δ ∼2.3 and ∼1.5 ppm. These clearly assignable signals demonstrate successful formation of both urethane and ester linkages in the polymer structure, providing a structural foundation for precise analysis of enzymatic degradation.

Complementary structural verification was achieved through Fourier-transform infrared (FTIR) spectroscopy (Fig. 1D). The spectrum displayed all expected characteristic absorptions: (i) a strong ester C=O stretch at 1725 cm^−1^, (ii) a broad urethane N–H stretch at 3320 cm^−1^, (iii) an amide II band (C–N stretch) at 1530 cm^−1^, and (iv) an ester C–O stretch at 1240 cm^−1^. These features showed excellent agreement with reference spectra of urethane and ester groups, supporting the expected chemical structure of the polymer.

To further validate the material’s properties, we characterized the thermal behavior of the synthesized PEA-PU using differential scanning calorimetry (DSC). The DSC results revealed that PEA-PU exhibits amorphous thermoplastic behavior, with a glass transition temperature (Tg) of approximately –17.6 °C, as determined by the half-height method (Supplementary Fig. 1). No melting transition was observed up to 200 °C. This thermal profile confirms the absence of crystalline domains, making the material particularly suitable for enzymatic degradation studies ^15,21^.

Taken together, these results demonstrate that PEA-PU is a structurally uniform, chemically well-defined polymer with straightforward synthesis and characterization. The material is optically transparent and exhibits distinct spectroscopic signatures, facilitating analytical tracking. More importantly, PEA-PU closely replicates the structural and chemical properties of post-consumer PU waste, making it a highly representative model system. These properties establish PEA-PU as an ideal model substrate for enzymatic degradation studies while also serving as a reproducible, standardized platform for developing robust PU degradation assays.

### 2.2. Enzymatic degradation of PEA-PU

Following comprehensive characterization of PEA-PU’s structure and physicochemical properties, we next evaluated its enzymatic degradability using two well-characterized polyester hydrolases: two well-characterized polyester hydrolases: ICCM (an engineered leaf compost cutinase variant) and BaCut1 (a cutinase from *Blastobotrys* sp. G-9). Although both enzymes demonstrate PU-degrading capability, prior studies have reported that ICCM exhibit greater polyester hydrolytic activity than BaCut1 under their respective optimal conditions^26,27^. We therefore employed PEA-PU as a model substrate to determine whether this well-documented activity difference could be reliably replicated, assessing its potential for standardized enzyme performance comparisons.

PEA-PU films were incubated with each enzyme for 36 hours under their optimal conditions (70 °C for ICCM, 37 °C for BaCut1). Enzymatic treatment induced significant morphological alterations in the films, which transitioned from an initially transparent state to an opaque and fragmented morphology (Fig. 2A). In contrast, control samples exhibited no structural or optical changes, demonstrating that the degradation was exclusively enzyme dependent.

**Figure 2:**
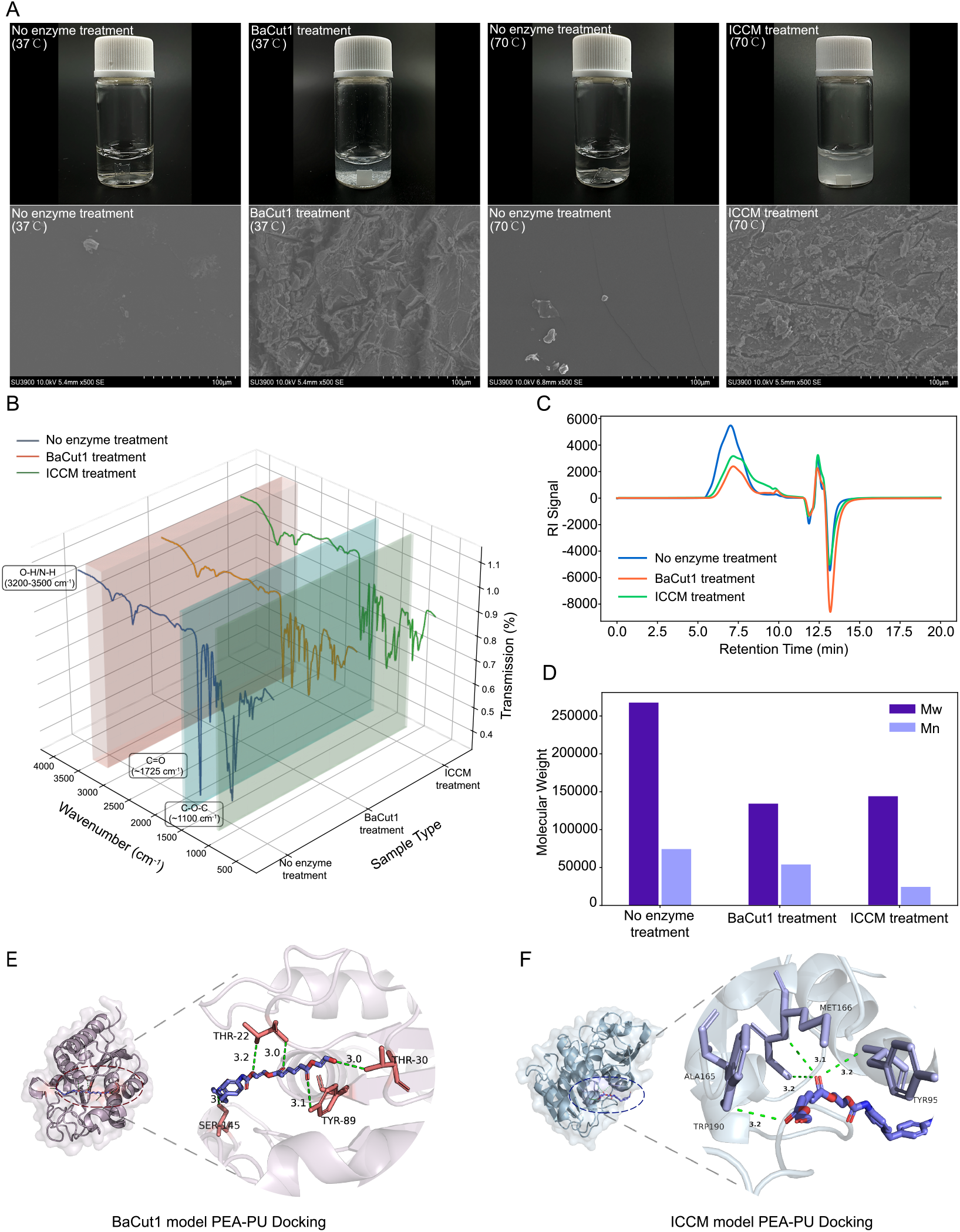
Enzymatic degradation of PEA-PU (A) Visual observation of film degradation & Surface morphology by SEM. PEA-PU films became turbid and fragmented after 36 h of incubation with ICCM (70 °C) or BaCut1 (37 °C), while enzyme-free controls remained optically transparent, indicating enzyme-specific hydrolytic activity. **(B) FTIR spectral changes following degradation. (C) GPC chromatograms**. GPC analysis was performed using DMF as the mobile phase after heat-assisted dissolution. **(D) Quantification of molecular weight reduction. (E, F) Molecular docking of enzyme–substrate complexes**. Docking simulations using AutoDock Vina showed that ICCM forms more hydrogen bonds with adipate ester linkages and binds substrates more deeply than BaCut1.

To examine surface-level enzymatic degradation, we analyzed PEA-PU films by scanning electron microscopy (SEM). While untreated films displayed smooth and uniform surfaces, enzyme-treated samples exhibited significant surface erosion (Fig. 2A). ICCM treatment induced pronounced degradation features, including deep pits and porous structures, whereas BaCut1 treatment caused milder surface disruption. These distinct morphological patterns reflect differential catalytic efficiencies between the two enzymes, consistent with previous reports^26,27^.

Further structural analysis by FTIR provided additional insights (Fig. 2B). Compared to untreated PEA-PU, enzyme-treated samples exhibited:(i) a broadened and slightly intensified O–H/N–H stretching band (3200– 3500 cm^−1^), suggesting the formation of hydroxyl and amine end-groups resulting from ester and urethane bond cleavage;(ii) a notable decrease in ester carbonyl (∼1725 cm^−1^) and C–O–C stretching (∼1100 cm^−1^) band intensities, confirming the breakdown of polyester segments.

To characterize the soluble degradation products, we performed chromatographic analysis of reaction supernatants. The PU monomer AA was quantified via liquid HPLC, while EG was detected by gas chromatography–mass spectrometry (GC–FID). Both monomers were exclusively present in enzyme-treated samples, with no detectable levels in negative controls. Notably, the observed AA:EG molar ratio (approximately 1:1) matched the theoretical stoichiometry of PEA-PU hydrolysis, wherein each ester bond cleavage generates equimolar AA and EG (Supplementary Table 1). This stoichiometric agreement establishes AA production as a robust quantitative indicator of EG release during enzymatic degradation.

Gel permeation chromatography (GPC) was then employed to quantify changes in polymer molecular weight following enzymatic treatment. Untreated PEA-PU exhibited a high molecular weight distribution, with a main retention time around 8.5 min. After enzymatic degradation, the distribution shifted toward higher retention times. Specifically, the main peak shifted to ∼10.0 min after ICCM treatment and to ∼9.2 min after BaCut1 treatment (Fig. 2C), consistent with the formation of lower-molecular-weight fragments. ICCM treatment led to a 67.9% reduction in number-average molecular weight (Mn) and a 47.3% reduction in weight-average molecular weight (Mw) (Fig. 2D), indicating substantial fragmentation into smaller oligomers. BaCut1 caused a relatively smaller but still significant decrease in Mn (∼28.2%) and Mw (∼48.5%). The corresponding polydispersity index (PDI) values are provided in Supplementary Figure S3. These molecular-level changes align with the distinct surface erosion patterns observed by SEM.

In addition, to understand how the enzymes interact with the PEA-PU, we performed molecular docking using representative PEA-PU oligomers. Our analysis revealed that both BaCut1 and ICCM preferentially bind to adipate ester linkages at their active sites (Fig. 2E–F), highlighting conserved features of ester bond recognition. Key interactions included hydrogen bonding networks and hydrophobic contacts near the cleavage site. Notably, ICCM exhibited stronger binding affinity than BaCut1, forming additional hydrogen bonds with scissile carbonyl oxygens and positioning the substrate deeper within its active site. These structural features— enhanced binding interactions and optimal substrate alignment—likely contribute to ICCM’s higher catalytic efficiency toward PEA-PU degradation compared to BaCut1.

Collectively, our integrated approach—combining microscopic, spectroscopic, chromatographic, and computational analyses—demonstrates that the chemically defined PEA-PU offers exceptional analytical tractability during enzymatic degradation. Upon enzymatic treatment, PEA-PU undergoes clear morphological changes—often visible to the naked eye—and generates well-defined soluble products. These monomers can be accurately identified and quantified using conventional chromatographic methods. Importantly, by assessing PEA-PU degradation with two established PU-degrading enzymes, we confirmed that this substrate reliably recapitulates their differential activities. The degradation profiles reflect enzyme-specific catalytic behaviors, enabling robust comparative analysis of PU-hydrolyzing activity. Given these attributes, PEA-PU emerges as an ideal model substrate for comparative enzyme characterization, providing a reliable basis for quantitative PU assay development.

### 2.3 Development of a coupled fluorescence assay for EG detection in PEA-PU degradation

The well-defined chemical composition, transparency, and homogeneity of PEA-PU enable direct observation and quantification of enzymatic degradation, with breakdown products readily analyzed by chromatography. However, like conventional substrates, PEA-PU has a critical limitation: while chromatographic analysis offers precise quantification, its low throughput hinders large-scale enzyme screening applications. To overcome this bottleneck, we designed an enzymatic cascade that converts the stoichiometric release of EG during PU hydrolysis into a sensitive fluorescent signal (Fig. 3A).

**Figure 3:**
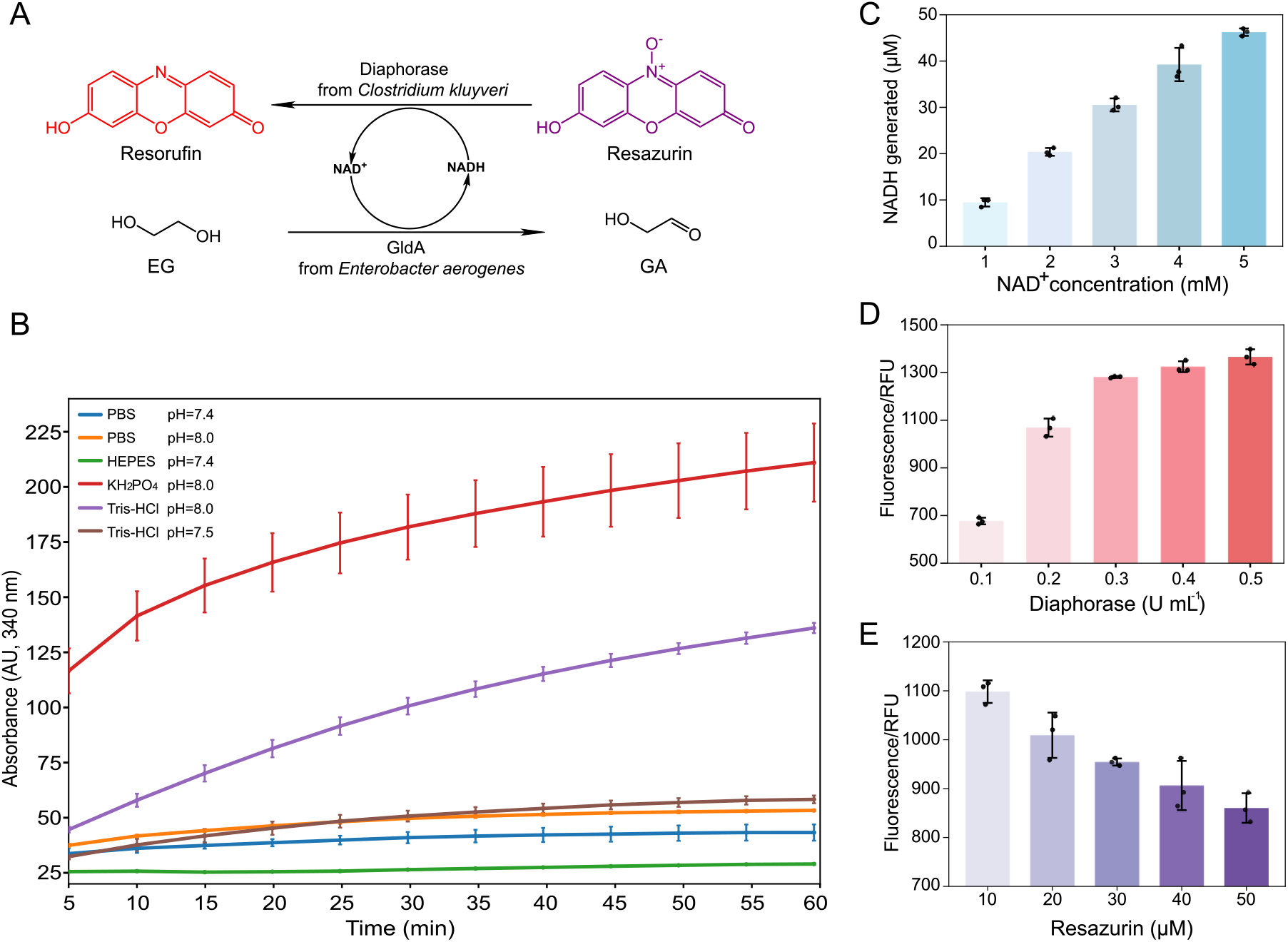
Development the coupled fluorescence assay. (A) Reaction scheme of the EG-coupled fluorescence assay. Schematic of the enzymatic cascade linking EG oxidation to resorufin fluorescence via GldA and diaphorase. **(B) Identification of the optimal buffer for GldA-mediated NADH generation. (C) Optimization of NAD**^**+**^ **concentration for maximal EG oxidation. (D) Optimization of diaphorase dosage for efficient signal conversion. (E) Selection of optimal resazurin concentration to avoid quenching**.

In this coupled system, we employed an NAD^+^-dependent glycerol dehydrogenase (GldA) from *Enterobacter aerogenes* ^28^ to oxidize EG to glyoxylic acid while reducing NAD^+^ to NADH. The resulting NADH is then used by diaphorase from *Clostridium kluyveri* ^29^ to reduce resazurin into resorufin, a highly fluorescent dye. This two-step enzymatic cascade ensures a direct, molecule-to-molecule correspondence between PU hydrolysis (EG release) and fluorescence intensity, enabling continuous and quantitative monitoring without extensive sample processing.

To establish robust performance of the coupled enzymatic fluorescence assay, we systematically optimized its reaction conditions. We began by evaluating buffer systems to maximize GldA activity. To avoid interference from downstream enzymes, GldA was assessed independently using EG as the substrate. NADH formation was monitored at 340 nm as a direct readout of enzymatic activity.

We tested six biological buffers: 50 mM Tris–HCl (pH 8.0 and 7.4), PBS (pH 8.0 and 7.4), HEPES (pH 7.4), and KH2PO4 (pH 8.0). Among these, KH2PO4 (pH 8.0) generated the highest NADH signal but exhibited poor reproducibility with significant inter-experimental variability. In contrast, 50 mM Tris–HCl (pH 8.0) supported consistently strong and stable NADH production with minimal variability (Fig. 3B). Additionally, Tris–HCl (pH 8.0) has been commonly used in prior studies involving diaphorase, supporting its suitability for the downstream coupling reaction. Based on these results, we selected 50 mM Tris–HCl (pH 8.0) as the standard buffer for all subsequent experiments.

Next, we determined the optimal concentration of the NAD^+^ cofactor. Increasing NAD^+^ from 1 mM to 3 mM accelerated NADH production, monitored via 340 nm absorbance (Fig. 3C). As NADH was produced in micromolar quantities, 3 mM NAD^+^ was sufficient to drive the reaction and yield a stable signal; further increases in NAD^+^ provided minimal additional benefit. Therefore, 3 mM was selected as the optimal concentration for subsequent assays (Fig. 3C).

We next optimized the concentration of diaphorase to ensure efficient signal transduction. Diaphorase activity was assessed by measuring fluorescence output using 20 μM NADH and 10 μM resazurin in 50 mM Tris–HCl (pH 8.0). Enzyme concentrations ranging from 0.1 U/mL to 0.5 U/mL were tested. After 30 minutes of incubation at room temperature, fluorescence (Ex/Em = 535/588 nm) increased with diaphorase concentration, plateauing at 0.3 U/mL. This suggests that 0.3 U/mL is sufficient for near-maximal signal output, with higher concentrations offering only marginal gains. Thus, 0.3 U/mL was selected as the optimal diaphorase concentration (Fig. 3D).

Finally, we evaluated resazurin, the fluorescent reporter, using 0.3 U/mL diaphorase and 20 μM NADH while varying resazurin (10–50 μM). Following the same incubation and detection protocol, fluorescence intensity decreased progressively with increasing resazurin concentrations. The highest signal was observed at 10 μM, suggesting that excess resazurin may induce self-quenching or inner-filter effects. Based on these results, 10 μM was selected as the standard resazurin concentration (Fig. 3E).

Through systematic optimization, we established a highly sensitive and reliable fluorescence-based assay for quantifying EG production from PU degradation. Under the finalized conditions (50 mM Tris–HCl, pH 8.0; 3 mM NAD^+^; 10 μM resazurin; 0.3 U/mL diaphorase), the assay delivers a strong, EG-dependent fluorescent signal across a broad detection range (0.25 – 25 mM EG), effectively capturing expected concentrations from polymer breakdown. The fluorescence generation rate (RFU) showed excellent linear correlation with EG concentration (R^2^ > 0.99 at all-time points from 30–120 minutes, measured at 10-minute intervals; Supplementary Table S2), demonstrating both precision and temporal stability. This assay enables continuous, high-throughput monitoring of PU degradation and is ideally suited for enzyme activity screening.

### 2.4 Validation of the coupled assay for PEA-PU degradation

To evaluate the quantitative performance of our coupled fluorescence assay, we first constructed calibration curves using EG standards ranging from 0.25 mM to 25 mM. Fluorescence generation rates (RFU/min), measured between 30 minutes and 120 minutes, exhibited a highly linear response (R^2^ = 0.9995; Figure 4A). We then applied this calibration to quantify EG released from PEA-PU films following enzymatic degradation by ICCM and BaCut1.

**Figure 4:**
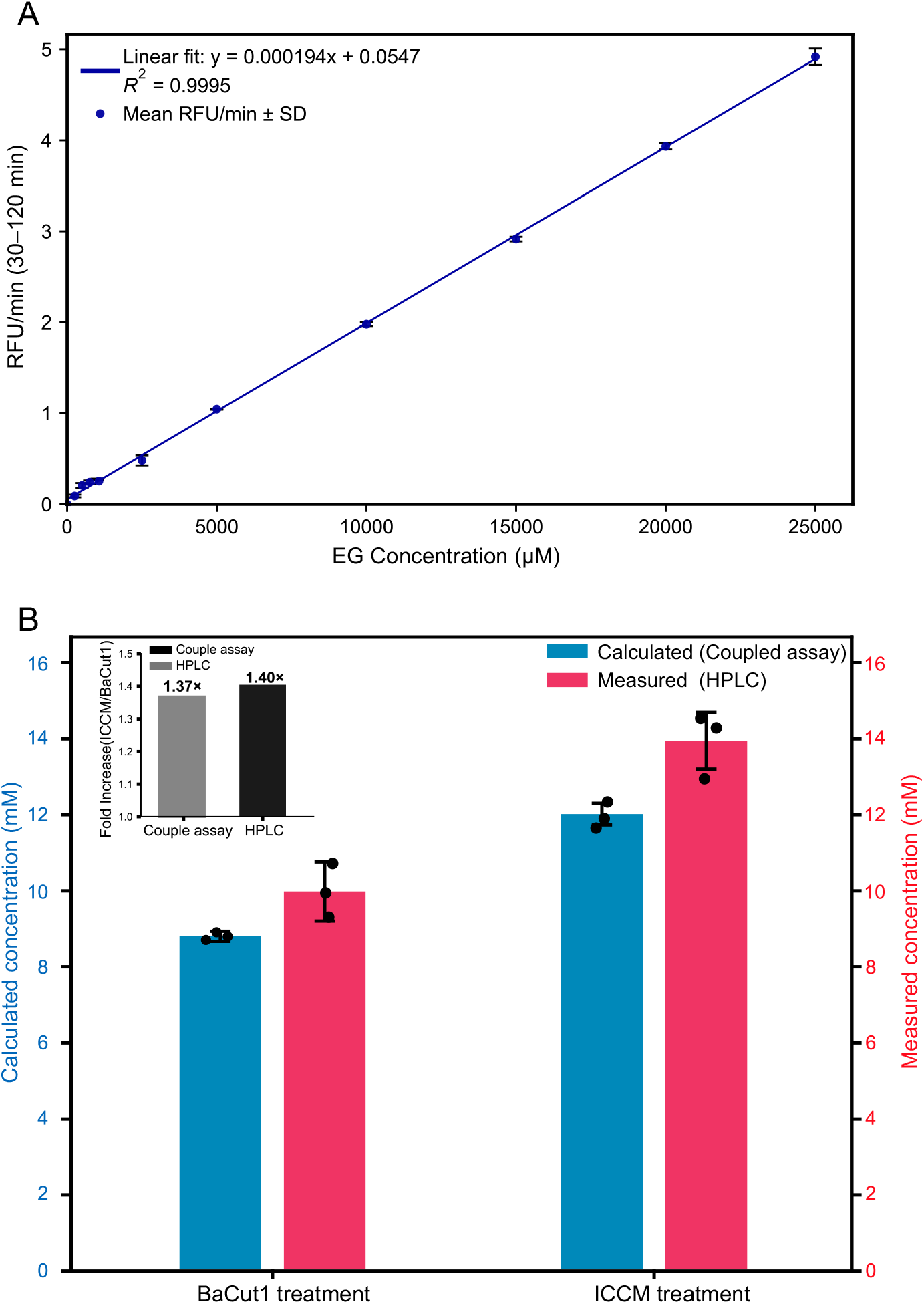
Validation of the coupled fluorescence assay for quantitative EG detection. (A) Standard curve for EG detection. Fluorescence rates (RFU/min) showed excellent linear correlation with EG concentration from 0.25 to 25 mM (R^2^ = 0.9995). **(B) Quantification of EG in enzymatic degradation samples**. Comparison of predicted EG concentrations (blue; derived from the fluorescence assay) and measured AA concentrations (magenta; determined by HPLC) in supernatants from PEA-PU films treated with BaCut1 and ICCM. Values reflect mean ± SD from three independent enzymatic reactions. **Inset**, Fold difference in enzymatic activity between ICCM and BaCut1 as determined by the fluorescence assay (1.37×) and HPLC (1.40×), demonstrating the assay’s ability to resolve enzyme-specific activity differences.

To independently validate these measurements, we used HPLC to quantify AA levels in the same samples, leveraging the known 1:1 stoichiometric co-release of EG and AA during PEA-PU hydrolysis (Supplementary Table 1). Because of this relationship, HPLC-measured AA levels serve as both a robust proxy for EG production (and thus overall polymer degradation) and a direct benchmark for assessing the accuracy of fluorescence-based EG detection. Notably, fluorescence-derived EG concentrations closely matched HPLC-measured AA values (Figure 4B), confirming the assay’s reliability for monitoring enzymatic degradation.

In addition to accuracy, our fluorescence assay also effectively resolved differences in enzymatic activity. As shown in Figure 4B (inset), the fluorescence assay detected a 1.37-fold greater EG release by ICCM compared to BaCut1. This ratio was highly consistent with the 1.40-fold difference measured via HPLC. This concordance underscores our fluorescence assay’s capacity to discriminate subtle variations in enzyme activity, a critical feature for HTS applications.

Finally, the fluorescence assay offers several practical advantages over HPLC, including reduced analysis time, lower sample volume requirements, and inherent compatibility with microplate-based high-throughput formats. These improvements address major limitations of conventional PU degradation assays and may accelerate the discovery and engineering of enhanced PU-degrading enzymes.

## 3. Conclusion

The development of effective HTS assays for PU-degrading enzymes is essential for discovering and optimizing biocatalysts to combat PU pollution. Here, we present an HTS assay that integrates a chemically defined PEA-PU substrate with an enzymatic cascade, converting stoichiometrically released EG into a sensitive fluorescent signal. Our assay shows excellent correlation with conventional analytical techniques (e.g., HPLC), demonstrating comparable reliability while offering the significant advantages of high-throughput compatibility, overcoming the limitations of traditional labor-intensive methods.

A key innovation of this work lies in our substrate design. Unlike proprietary substrates such as Impranil DLN (with undefined composition and obscure degradation pathways), our PEA-PU system establishes clear structure-activity relationships, directly linking specific polymer bonds to their degradation products (AA and EG). This mechanistic clarity enables a deeper investigation of enzymatic mechanisms, supporting rational enzyme engineering and advanced screening strategies.

Beyond its fundamental advantages, the assay offers practical benefits, including rapid processing, minimal sample requirements, and seamless compatibility with multi-well plate formats. These features significantly reduce time and resource demands, enabling efficient screening of environmental isolations and large mutant libraries. Such capabilities make our assay particularly valuable for large-scale applications, from functional metagenomic discovery to directed evolution campaigns.

Future refinements could further enhance the assay’s versatility, such as extending its sensitivity to etherbased PU cleavage markers or incorporating alternative fluorophores, enzymatic cascades, and detection modalities. Integration with emerging technologies—such as artificial intelligence–assisted enzyme design, automated liquid handling, and ultra-high-throughput microfluidics— could further enhance screening efficiency and success rates.

Ultimately, our HTS assay for PEA-PU degradation provides a robust, scalable, and mechanistically precise tool for accelerating the discovery and engineering of PU-degrading enzymes. By utilizing a structurally realistic substrate, the assay ensures that identified hits exhibit industrially relevant depolymerization activity, effectively bridging the gap between laboratory discovery and real-world PU waste degradation. This approach holds significant promise for advancing sustainable PU waste management, a critical step toward achieving a circular plastics economy.

## Acknowledgements

We acknowledge the support from the Biosciences Central Research Facility (BioCRF (GZ)) and the Materials Characterization and Preparation Facility (MCPF (GZ)), both of which are central research facilities of the Hong Kong University of Science and Technology (Guangzhou) (HKUST(GZ)). Their assistance in experimental procedures and material characterization was essential to this study.

## 5. Materials and Methods

### 5.1. Materials

Unless otherwise stated, all reagents were of analytical grade and used as supplied. Hydroxyl-terminated poly (ethylene adipate) (PEA, Mn ≈ 1000 g/mol, CAS 24938-37-2), 4,4′-methylenebis (phenyl isocyanate) (MDI, CAS 101-68-8), and anhydrous N, N-dimethylformamide (DMF, CAS 68-12-2) were purchased from Aladdin Industrial Corporation (Shanghai, China).

Resazurin (≥98%, CAS No. 550-82-3) and Diaphorase from *Clostridium kluyveri* (3.0-20.0 units/mg protein, CAS No. 9001-18-7) were purchased from Sigma-Aldrich (St. Louis, MO, USA). β-Nicotinamide adenine dinucleotide (NAD^+^, >98%, CAS No. 53-84-9) was obtained from Roche Diagnostics GmbH (Mannheim, Germany).

The polyurethane-degrading enzymes ICCM and BaCut1 were constructed as described: ICCM based on engineered LCC variants (F243I/D238C/S283C/N246M) per Nature 2022 (DOI:10.1038/s41586-022-04599-z), and BaCut1 based on the reported sequence from *Blastobotrys* sp. G-9 (DOI:10.1016/j.jhazmat.2024.134493), was codon-optimized, synthesized by Tsingke Biotechnology Co., Ltd. (Beijing, China), and cloned into the pET32a expression vector. Glycerol dehydrogenase (GldA) from *Enterobacter aerogenes* was recombinantly expressed in-house in *E. coli* BL21(DE3) using a pET32-based vector and purified by Ni–NTA affinity chromatography as described in Section 5.2.3.

All other routine chemicals, buffer components (e.g., Tris base, NaCl, imidazole), and consumables were purchased from Sigma-Aldrich unless otherwise stated. Ultrapure water (18.2 MΩ·cm) was prepared using a Milli-Q® Integral water purification system (Merck Millipore, USA). Black, flat-bottom, low-volume 96-well plates were obtained from Corning® (Corning, NY, USA) for fluorescence measurements.

### 5.2. Methods

#### 5.2.1. Synthesis of PEA-PU Films

PEA was vacuum dried at 120 °C for 1 hour prior to use. In a nitrogen-purged three-neck flask, equimolar amounts of PEA and MDI (NCO: OH = 1:1) were dissolved in anhydrous DMF (10% w/v). The reaction mixture was stirred at 70 °C for 1 hour under a dry nitrogen atmosphere. The resulting viscous prepolymer was cast into a PTFE mold and vacuum-dried at 60 °C for 24 hours, yielding uniform transparent films of approximately 0.2 mm thickness. Films were stored in a desiccator prior to use.

#### 5.2.2. Structural and Thermal Characterization

^1^H NMR spectra were recorded on a Bruker Avance III 400 MHz spectrometer using DMSO-d_6_ as solvent. Chemical shifts were referenced to the residual solvent peak at δ 2.50 ppm.

ATR-FTIR spectra were obtained using a Bruker Vertex 70V FTIR spectrometer coupled with a Hyperion II microscope (Bruker, Germany), operated at 4 cm^−1^ resolution with 32 scans per sample.

Differential scanning calorimetry (DSC) measurements were performed on a DSC 2500 (TA Instruments, New Castle, DE, USA). Approximately 5 mg of each PEA-PU film was sealed in an aluminium Tzero™ pan with a matching lid. Samples were equilibrated at –50 °C for 1 min, heated to 200 °C at 10 °C min^−1^, held for 1 min, cooled to –50 °C at the same rate, and then reheated to 200 °C. Thermograms from the second heating scan were used for analysis to remove thermal-history effects. The glass-transition temperature (Tg) was determined from the midpoint of the heat-capacity step in the second heating trace.

SEM analysis was conducted to assess surface morphology before and after enzymatic degradation. Samples were sputter-coated with a thin layer of platinum to ensure conductivity. Imaging was performed using a Hitachi SU3900 Scanning Electron Microscope at an accelerating voltage of 10.0 kV, a working distance of 5.4 mm, and a magnification of ×500 under secondary electron (SE) mode. Scale bars were set to 100 μm.

#### 5.2.3. Protein Expression and Purification

The enzymes GldA, ICCM, and BaCut1 were recombinantly expressed and purified for this study. The corresponding genes for each enzyme were cloned into the pET32a(+) expression vector, which incorporates an N-terminal His_6_-tag for affinity purification. The resulting plasmids were transformed into E. coli BL21 (DE3) host cells.

For protein expression, cultures were grown in Luria-Bertani (LB) medium containing 100 μg/mL ampicillin at 37 °C with vigorous shaking until the optical density at 600 nm (OD_600_) reached 0.6–0.8. Protein synthesis was then induced with a final concentration of 0.2 mM isopropyl β-D-1-thiogalactopyranoside (IPTG), and the cultures were further incubated at 16 °C for 16–20 hours.

After incubation, cells were harvested by centrifugation (4,000 × g, 30 min, 4 °C), and the cell pellet was resuspended in lysis buffer (50 mM Tris-HCl, pH 8.0, 150 mM NaCl). The cells were disrupted by sonication on ice, and the crude lysate was clarified by high-speed centrifugation (30,000 × g, 30 min, 4 °C) to remove cell debris.

The clarified supernatant was loaded onto a Ni Sepharose 6 Fast Flow affinity column (Cytiva) pre-equilibrated with the lysis buffer. The column was washed with the same buffer containing 20 mM imidazole to remove non-specifically bound proteins. The target His-tagged protein was subsequently eluted using a buffer containing 300 mM imidazole. The eluted fractions were desalted and concentrated using Amicon® Ultra-15 centrifugal filters (10 kDa MWCO, Merck Millipore). The purity of the final protein samples was confirmed by SDS-PAGE, and their concentrations were determined spectrophotometrically at 280 nm using a NanoDrop™ 2000 spectrophotometer (Thermo Fisher Scientific).

#### 5.2.4. Enzymatic Degradation of PEA-PU Films

PEA-PU films were cut into 0.8 cm × 0.8 cm squares. Each film sample was incubated in 1.0 mL of 50 mM Tris–HCl buffer (pH 8.0) containing ICCM (13.95 μg/mL) or BaCut1 (125 μg/mL) at their respective optimal temperatures (70 °C and 37 °C) for 36 hours. Control samples lacked enzyme. Reaction supernatants were collected for product analysis.

#### 5.2.5. Chromatographic Analysis of Degradation Products

To validate the enzymatic degradation of PEA-PU films, the resulting hydrolysis products were detected using gas chromatography and high-performance liquid chromatography.

##### Gas Chromatography–Flame Ionization Detection (GC-FID)

Ethylene glycol (EG) in the enzymatic degradation supernatants was analyzed by a third-party analytical laboratory using a Shimadzu GC system equipped with a flame ionization detector (FID). EG was detected at a retention time of approximately 4.73 minutes. Quantification was based on an external standard calibration curve (5–20 ppm), which showed excellent linearity (R^2^ > 0.998).

##### High-Performance Liquid Chromatography (HPLC)

Adipic acid (AA) was analyzed using a Thermo Scientific Vanquish Core HPLC system (Thermo Fisher Scientific, Germany) equipped with a Shim-pack GIST C18 AQ column (3 μm, 150 mm × 4.6 mm; Shimadzu, Japan). The mobile phase consisted of 0.1% trifluoroacetic acid (TFA) in water (solvent A) and acetonitrile (solvent B). Gradient elution was performed as follows: 0–20 min, 5–95% B; 20–24 min, held at 95% B; 24– 25 min, ramped back to 5% B; followed by a 5-minute re-equilibration at 5% B. The flow rate was 1.0 mL/min, the detection wavelength was set at 210 nm, and the injection volume was 10 μL. Quantification was based on an external calibration curve (1–100 ppm), which showed excellent linearity (R^2^ > 0.999).

#### 5.2.6. Coupled Fluorescence Assay for EG Detection

A fluorescence-based coupled enzyme assay was developed to quantify ethylene glycol (EG) released from PEA-PU degradation. Reactions were performed in black 96-well plates with a final volume of 200 μL. Each reaction contained 50 mM Tris-HCl buffer (pH 8.0), 3 mM NAD^+^, 10 μM resazurin, 0.3 U/mL diaphorase, 0.25 mg/mL GldA, and either EG standards or samples.

To construct the standard curve, EG standards (0.25–25 mM) were prepared in assay buffer and incubated at room temperature. Fluorescence was monitored at Ex/Em = 535/588 nm using a Thermo Scientific Varioskan™ LUX microplate reader in kinetic mode for 180 minutes, with readings collected every 5 minutes. The fluorescence rate (RFU/min) was plotted against EG concentration, yielding a linear regression with R^2^ ≥ 0.9995.

For experimental samples, 20 μL of degradation supernatant was added to 180 μL of assay buffer. Fluorescence was recorded under identical conditions, and EG concentrations were interpolated from the standard curve. All measurements were performed in triplicate.

